# Two common issues in synchronized multimodal recordings with EEG: Jitter and Latency

**DOI:** 10.1101/2022.11.30.518625

**Authors:** Seitaro Iwama, Mitsuaki Takemi, Ryo Eguchi, Ryotaro Hirose, Masumi Morishige, Junichi Ushiba

## Abstract

Multimodal recording using electroencephalogram (EEG) and other biological signals (e.g., electromyograms, eye movement, pupil information, or limb kinematics) is ubiquitous in human neuroscience research. However, the precise time alignment of data from heterogeneous sources is limited due to variable recording parameters of commercially available research devices and experimental setups. Here, we introduced the versatility of a Lab Streaming Layer (LSL)-based application for multimodal recordings of high-density EEG and other devices such as eye trackers or hand kinematics. To introduce the benefit of recording multiple devices in a time-synchronized manner, we discuss two common issues in measuring multimodal data: jitter and latency. The LSL-based system can be used for research on precise time-alignment of datasets, such as detecting stimulus-induced transient neural responses and testing hypotheses well-formulated in time by leveraging the millisecond time resolution of the system.

## Introduction

Scalp electroencephalogram (EEG) non-invasively records data from human neural systems implicated in neural processing underlying cognitive and sensorimotor behavior (Cavanagh and Frank 2014; Lachaux et al. 1999; Makeig et al. 2004; Michel and Koenig 2018; Pfurtscheller and Lopes Da Silva 1999). Although the quality of measured signals is affected by various factors (e.g., volume conduction, noise, and artifacts), EEG is widely used because of its cost-effectiveness and convenience. Scalp EEG signals provide limited information from approximately 10^7^ neurons (Buzsáki et al. 2012; Sejnowski et al. 2014); therefore, it is difficult to identify the functional representations of neural structures based on EEG signals alone. The complexity of analyzing neural processing on EEG can be compared to a journalist reporting from a stadium (Biasiucci et al. 2019, Figure 1A). Although invasive techniques can measure spike activities or local field potentials by entering the stadium and interviewing the audience of the event, EEG solely measures the sound of the crowd from outside. Therefore, the event responsible for the measured signals (Figure 1B) may be identified by using other modalities that record data streams containing behavioral (e.g., electromyogram (EMG) or kinematics) or physiological (e.g., pupil diameter or skin conductance change) information. These data elucidate the consequences of neural computation and reflect its difference among the experimental conditions (i.e., what events happened in the stadium).

**Figure 1.**
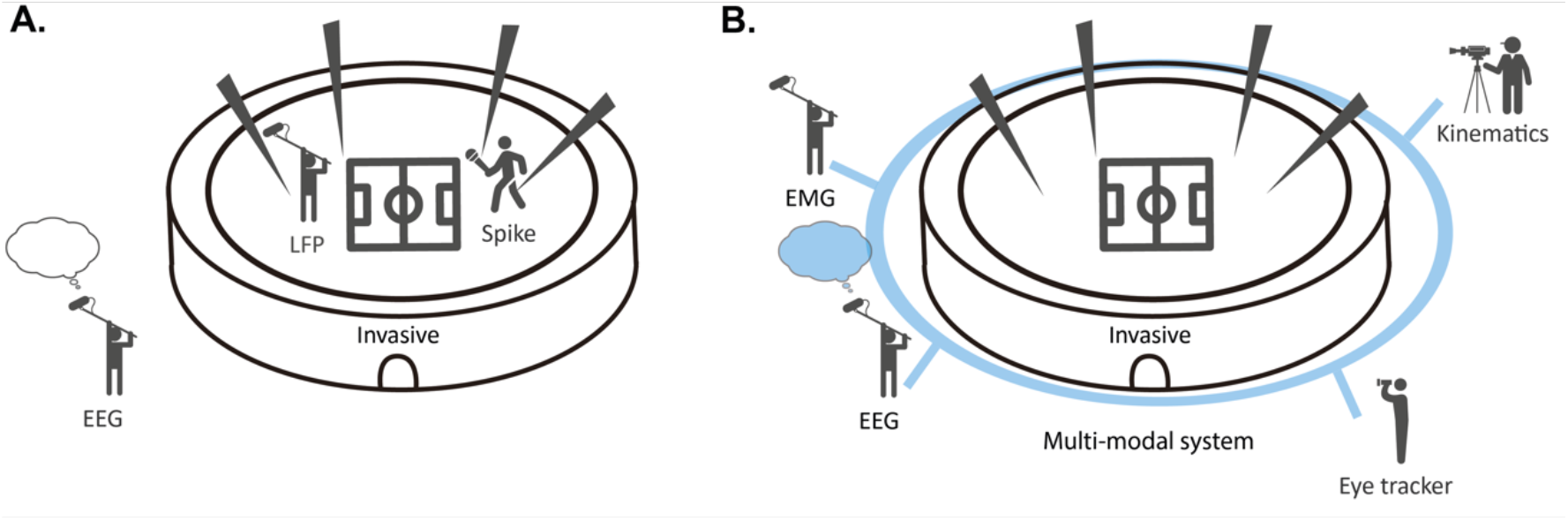
Schema of neural signal measurements. **(A)** An event of interest taking place in a stadium represents the recording of neural data. Direct access inside the stadium is only allowed for invasive recordings, indicating the direct placement of electrodes on and/or into the brain. Scalp EEG signals provide an indirect measurement of the summed activity of neurons (audience in the stadium). **(B)** Multimodal recording of biosignals helps to identify the neural representation. Recording of peripheral activities, such as EMG signals, is analogous to the interview of people receiving broadcast signals. Eye tracking and motion capture support the identification of the neural representation along with the EEG signals.

Multimodal recording of biological responses, such as combining data from the EEG amplifier and sensor devices, is a promising approach to exploring behaviorally relevant neural processing (Gilzenrat et al. 2010; Stangl et al. 2020). Studies on perception frequently combine the EEG setup and eye trackers to examine the internal processing of given stimuli (Lee et al. 2010; Plöchl et al. 2012; Simola et al. 2015). In addition, scalp EEG is often used to assess the neurological symptoms that modulate synaptic properties of sensorimotor pathways (Abbruzzese and Berardelli 2003; Cruccu et al. 2008; Ray et al. 2020; Shibasaki et al. 1986; Tangwiriyasakul et al. 2014). The latency of evoked potentials is used to diagnose patients by analyzing EEG data and peripheral recordings aligned with the timing of stimulus presentation (Tscherpel et al. 2020).

Therefore, determining the timing of stimulus presentation that elicits event-related responses requires precise alignment of multimodal data and EEG signals to ensure consistent temporal properties across trials.

However, there is not sufficient discussion about how inadequate data synchronization affects the neurophysiological analysis of multimodal dataset in neuroscience studies. A variety of recording parameters of each device, such as sampling rate, protocols of data transfer, and hardware, affect the multimodal measurements. While recent commercial research devices have communication protocols to connect to others, the protocols are variable. Both digital and analog communication protocols are commonly used to send data packets to a host computer, for instance via a local network, acquiring data with external analog-digital converters, or transistor-transistor logic (TTL) based event timing recording for post-hoc segmentation. Moreover, due to the temporal variance of data acquisition resulting from the heterogeneity of device settings, multimodal data acquisition requires additional costs to harmonize data from different sources. One metric to assess the efficacy of data acquisition is latency, which is the difference between the timings of actual data measurement and reception by the computer. Although post-hoc alignment can correct latency that was constant throughout the experiment, additional efforts are needed to correct latency that was variable during measurements (jitter). The precise analysis of multimodal data requires the common time stamps with which those data are aligned.

Here, we evaluated the effects of jitter and latency on acquisition of multimodal and EEG signals and introduced an experimental setup called Lab Streaming Layer (LSL). LSL improves the synchronization of multimodal and EEG signals without sharing hardware triggers, thereby enabling unified data collection from different sources (Stenner et al., 2022). We evaluated the efficacy of the LSL-based system and compared it to a typical experiment setup, namely event-related potential (ERP) acquisition. Because the implementation of the software-based time synchronization with the LSL-based system is relatively complex, the software used in the present paper was made publicly available as open-source software with step-by-step documentation. Then, we compared multiple experimental setups in terms of several perspectives, including time-alignment, extension capability, and data collection. This software system would contribute to constructing a reliable experimental setup for researchers using EEG to test their hypotheses.

### Experimental setup for a multimodal recording of human data

In human neuroscience experiments, multimodal recording of biosignals is generally achieved by the simultaneous use of multiple commercial devices. Several methods are available to synchronize the signals from EEG and other devices (Figure 2A), such as auxiliary channels implemented in the EEG amplifier, hardware-based triggers generated by external devices, and software-based synchronization using the LSL system.

**Figure 2.**
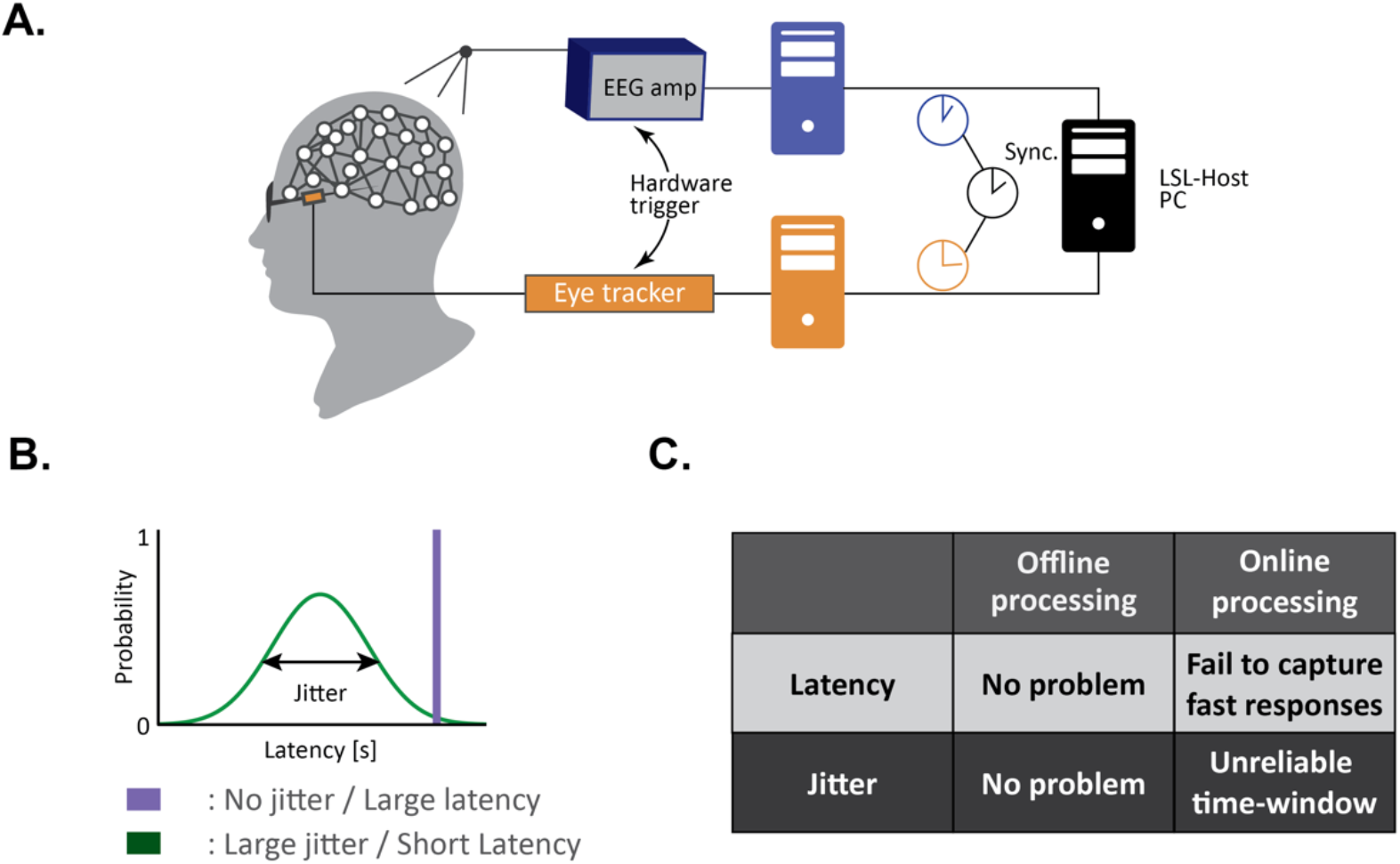
Synchronized multimodal recordings with EEG. **(A)** EEG recording setup for multimodal data acquisition. Hardware trigger represents a simultaneous input delivered to two individual devices. In the LSL setup, the host computer aligns two received signals with the common timestamps. **(B)** A schematic of jitter and latency relationships. In the no-jitter large-latency case represented by a purple bar, the latency of samples from a device is fixed. In contrast, in the large-jitter short-latency case, latency for online processing becomes variable. **(C)** Summary of effects of jitter and latency on online and offline processing. In offline processing, post-hoc processing can correct the latency and jitter effects as long as the data were collected at a constant sampling rate while they respectively induce effects on online processing.

#### Auxiliary channels implemented in the EEG amplifier

Recent EEG amplifiers provide auxiliary inputs to simultaneously acquire other electrical biosignals, such as EMG, electrocardiogram, and electrooculogram. This functionality enables the simultaneous acquisition of multimodal signals with no latency or jitter difference with EEG because amplifiers send data packets that contain all the acquired signals to computers. Signal processing can be improved by investigating the precise temporal profiles of neural responses by linking the acquired multiple signals (Al et al. 2020; Richter et al. 2017). However, the signal variety is limited by the specifications of the amplifiers.

#### Hardware-based triggers generated by external devices

External devices, such as eye or motion trackers and stimulation presenters, are regularly used for perception, cognition, and sensorimotor studies (Lee et al. 2010; Plöchl et al. 2012; Simola et al. 2015). Although a variety of biosignals can be used to indicate internal neural processing involved in a specific task (e.g., pupil diameters, galvanic skin response, and oxygen saturation level), the precision of time alignment depends on the availability of hardware-based triggers, such as transistor-transistor logic, serial communication, or analog output. In the absence of such triggers, the precise alignment of the multimodal dataset is challenging.

#### Software-based time synchronization system

An alternative approach for the time alignment of signals from multiple devices is software-based synchronization, which aligns the acquired signals with sub-millisecond accuracy using a common time series with the clock of the computer as a reference. Human neuroscience studies already use versatile platforms that facilitate temporal alignment, e.g., openly available software: OpenViBE (Renard et al. 2010), Naturalistic Experiment Design Environment (Jangraw et al., 2014), Unified Suite for Experiments (Marcus et al., 2019), OpenSync (Razavi et al., 2022) and LSL (Stenner et al., 2022); proprietary software: iMotions or Presentation. In particular, the recent open software OpenSync enables the experimenter to flexibly design the experimental setup combined with other platforms such as Unity. Such software can construct a local network to accumulate signals from connected client computers with time stamps and automatically align them. Software-based integration of multiple sources of biosignals enhances the flexibility of the experimental setup and precision of synchronization.

#### Jitter and latency

If more than two devices are used in an experimental setup, it is necessary to evaluate the precision of time synchronization from two perspectives: jitter and latency. Jitter indicates the timing variability when a computer receives signals, and latency indicates the absolute time for samples to reach computers (Figure 2B). Signals recorded by the devices reach computers after a latency. Here, we considered two protocols to send the acquired data packets to a computer: no-jitter and large-latency, and large-jitter and short-latency. In the former case, computers receive signals after a constant delay because the device sends packet data after the specified number of samples is accumulated. However, in the latter case, computers receive signals after a variable delay because the device sends packet data when the computer requests it. Although the latter method decreases the average latency, it causes a jitter.

Jitter and latency have different effects on data acquisition for offline and online processing (See also a summary matrix: Figure 2C). For offline data collection, both latency and jitter do not significantly impact data analysis if all modalities are time aligned and samples from each device are measured at a constant and accurate sampling rate. For instance, the hardware triggers used in the typical experiment setup can achieve the time synchronization of multiple devices by providing triggers at the timing of events such as stimulus presentation. However, if one of the devices used in an experimental setup does not generate continuous time point data, the accuracy of post-hoc time alignment of multiple modalities is dependent on the jitter and latency of the device. For instance, if the hardware triggers are received by an analog-to-digital converter that only detects the timing of signal input (i.e., only records the timing of trigger arrival), there is no information to correct the jitter and latency of signals from other devices during post-hoc analysis.

For online data collection, jitter induces the variability of available latest data, and latency induces the absolute delay of available signals. In zero-jitter and large-latency cases, the online processed signals used to deliver triggers to other devices or sensory stimuli to participants contingent on processed signals have a fixed delay specified by latency. If the latency is too significant to capture the neural responses of interest, this experimental setup will fail to capture the state-dependent difference in the response to external stimuli.

In contrast (large-jitter and short-latency cases), the online processing would use the latest samples while the exact timepoints available at each routine vary due to the jitter. When the online analysis of biosignals runs at a specific interval, the experimenter needs to admit the variance of available sample size at each process. The effect becomes more significant in the experimental paradigm that triggers external devices contingent on the biosignals. Because the stimulation timing is determined on the basis of the latest signals, the variable latency between signal acquisition and stimulation presentation can be a confounding variable during analysis (Tomasevic et al., 2017).

These two examples have pros and cons for the temporal precision and efficacy of online biosignal processing. If the brain-state-dependent paradigm, such as EEG-dependent stimulation techniques, requires a small time window of interest to capture characteristics of transient neural oscillations, the large-jitter short-latency case would be the better choice. However, the jitter of feedback signals would decrease the learning rate of neurofeedback training due to the variability of signals used for online feedback. Hence, in the case of the neurofeedback paradigm, the decrease in jitter would be beneficial for the controllability of targeted neural signaling by admitting a fixed window of latency (Belinskaia et al., 2020). As EEG studies leverage time-resolved features of the acquired signals in the order of milliseconds (e.g., amplitude profile and phase synchronization), data acquisition timing is critical to capture the phenomena of interest. The following section discusses the versatile experimental system that achieves data synchronization with sub-millisecond accuracy across multiple devices with long-term stability.

### Simulation analysis of jitter-induced changes in signal-to-noise ratio

We tested the effects of time synchronization accuracy by software-based synchronization in terms of signal-to-noise ratio (SNR) between two types of ERPs, namely somatosensory-evoked potentials and visual-evoked potentials (VEPs). Using the LSL-based experimental setup shown in Figure 3A and a data acquisition system, we simultaneously measured EEG, EMG, eye movements, and stimulus timing (Time course of preprocessed signals from a typical trial: Figure 3B). A checkerboard presented to the participants was inverted to elicit the VEPs (i.e., stimulus presentation). Changes in pupil diameters were concurrently observed at the time of stimulation presentation (Liao et al. 2016; Wang and Munoz 2014). Note that the preliminary demonstration is to empirically show the significance of jitter for typical EEG analysis.

**Figure 3.**
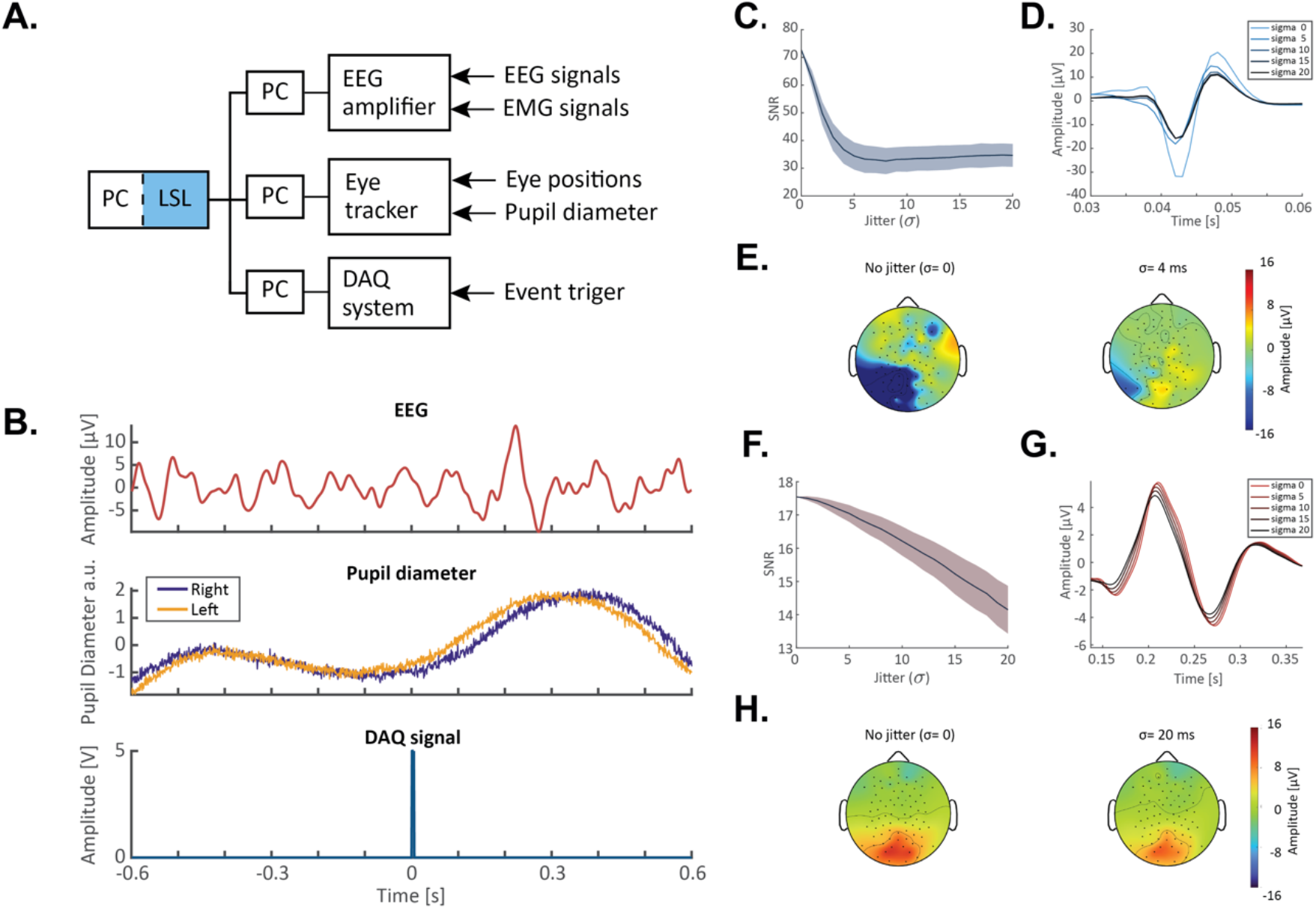
Empirical demonstration of LSL-based time synchronization. **(A)** Experimental setup used for evaluation. **(B)** Time course of preprocessed signals of multimodal data from a typical trial aligned by the LSL-based application. Representative EEG signals from the Oz channel are shown at the top. The pupil diameter exhibits its modulation triggered by visual stimulation (checkerboard) presentation when the DAQ system received a 5V pulse. **(C)** Changes in signal-to-noise ratio (SNR) due to artificial jitter augmentation for SEP data. Shade represents 1 standard deviation computed based on SNR calculated at 1000 times for each condition, reflecting the variance due to the random sampling of artificial jitter. **(D)** Time-courses of N20 potentials derived from jittered data. **(E)** Topographic representations of the grand average SEP derived from the no jitter and jittered data. **(F)** Changes in signal-to-noise ratio (SNR) due to artificial jitter augmentation for VEP data. **(G)** Time-courses of P3 potentials derived from jittered data. **(E)** Topographic representations of the grand average VEP derived from the no jitter and jittered data.

We evaluated the negative (N20) and positive (P150) peaks observed after somatosensory and visual stimulation, respectively. These components are prominent peaks elicited by sensory stimuli and are frequently used in basic neuroscience studies (Stephani et al. 2020, 2021) and clinical investigations (Halliday et al., 1973). For instance, the latency and amplitude of these responses are used to probe aging and evaluate neurological symptoms because they are linked to the transsynaptic pathways (Betsuin et al. 2001; Teyler et al. 2005). We recorded the stimulus-evoked EEG signals from a single participant during somatosensory and visual stimulations using a 128-channel HydroCel Geodesic Sensor Net (GES 400; Electrical Geodesics, Inc., Eugene, OR, USA). The participant provided written informed consent, and the study design was approved by the local Ethics Committee of Keio University (approval number: 2021-118). The experiment was performed in accordance with the Declaration of Helsinki. We used the Cz channel as the reference and acquired EEG signals were preprocessed with standard bandpass (20-200 Hz for SEP and 0.1-45 for VEP) and notch (50 Hz) filtering with a third-order Butterworth filter. The top panel of Figure 3B displays the evoked potentials from the electrode placed over the visual cortex (Oz). We presented the visual and somatosensory stimuli 300 times each and measured the EEG, pupil diameter (Tobii pro spectrum, Tobii Technology, Inc, Stockholm, Sweden), and stimulation timing using a digital-analog converter (USB-63 343, NI, Inc, Austin, TX, USA).

To assess the impact of jitter on the quality of ERPs, simulations were performed by randomly adding a variety of jitters to each segment. Then, we extracted ERPs by averaging the segmented trials at the sensor-space. The jitter sizes were sampled from a Gaussian distribution with mean 0 and standard deviation from 0 to 20 ms, respectively. SNR values of ERPs were computed using the baseline period as reference (amplitude of EEG during the last 1s before the stimulus presentation). Because characteristic evoked responses were originally observed at the time point of interest (20 ms for SEP and 150 ms for VEP), the attenuation of such responses should be attributed to the compromised quality of signal alignment. The simulation results indicated that even slight jitter (~2ms) induces a decline in SNR due to the unreliable trigger signals (Effects of jitter on the SNR of SEP: Figure 3C). The waveforms were gradually attenuated by the jitter (Changes in the SEP waveform in each jitter condition: Figure 3D). As we used LSL-based time synchronization during measurement, the offline segmented data exhibited representative waveforms of ERP and spatial extent in keeping with their purported sources (Figure 3E, left panel). However, the spatial extent of ERP is also affected due to jitter (Topographic representation of SEP: Figure 3E, right panel). By adding random jitter with a 4 ms standard deviation, the negative peak of the N20 component localized around the left parieto-temporal region was diminished (Figure 3E, left panel). An identical analysis was conducted for the occipital region (Oz) during VEP acquisition (Figure 3F-H). The decline in SNR is robustly observed for the different ERPs, although the effect size is small because of the relatively slower dynamics of the VEP components compared to that of SEP

## Discussion

### Advantages and disadvantages of experimental setups

Temporal precision during recording is paramount for multimodal recording. In addition, flexibility of the experimental setup and the convenience of using and sharing data are important for the versatility of the experimental setups.

In the present study, we compared hardware triggers and software-based time synchronization methods including LSL as the system for multimodal recordings from diverse perspectives: latency, jitter, extension capability, installation cost, time alignment, and data sharing.

The experimental setup combined with hardware-based triggers generated by external devices may have longer latency and larger jitter than the LSL-based system because different computers independently receive the data, each with its distinct clock. Moreover, jitter increases when individual computers receive data at different timings. Meanwhile, the LSL-based system does not have such problems because signals of the amplifier and auxiliary input are synchronously acquired and are aligned by the host computer using the LSL-based system.

The extension capability of the LSL-based system is relatively higher than that of the auxiliary inputs of EEG amplifiers or hardware since the LSL system can receive data from multiple devices after adding the corresponding modules. However, implementation of the LSL system can be technically challenging because it requires the modules for each device and the host computer, while the use of the existing applications for each product allows easy data acquisition. To address this issue, we have published the software for the LSL system-based recorder and data acquisition programs that support the devices frequently used in human neuroscience studies (see Data Availability Statement).

Time alignment and data sharing by the LSL-based systems are more manageable than saving data from each device as individual files because the system saves an integrated dataset such as a single file as Extensible Data Format (Extensible Data Format, 2015), which temporally aligns the data channels. In contrast, individualized datasets require post-hoc integration using custom scripts.

### Efficient ERP acquisition using the software-based time synchronization

The present paper highlighted the importance of precise time alignment of multimodal data. The prototype demonstration shown here was the acquisition of event-related responses and the comparison of SNR changes of ERPs caused by artificial jitter. Time synchronization by the LSL-based platform elicited higher SNR without jitter augmentation, which segmented the acquired EEG data based on asynchronously measured stimulus triggers. Because experimental setups may simultaneously use an EEG amplifier and external devices, the trigger imitates the signal of biological responses measured by external devices. While the topographic representation derived from the no jitter condition exhibited a typical spatial pattern, the pattern was impaired by adding a relatively small jitter (e.g., 4 ms) which exhibited the half SNR of N20 components. Therefore, the precision of trigger timing affects the stimulus-locked responses, such as ERPs. The empirical preliminary demonstration suggests a common software-based platform for multi-modal data acquisition from heterogeneous devices available as an open-source or commercial application is indispensable for analyzing EEG signals combined with other modalities without hardware triggers. However, further systematic comparison of multiple software platforms is warranted to determine the optimal procedure to handle the jitter and latency problems.

### Implications of the software-based time synchronization system for physiological experiments

The time-synchronized acquisition of multimodal datasets enhances the robustness of experiment results and facilitates the identification of correlations among modalities. Although accurate temporal synchronization depends on the experimental design, reduced jitter and latency in data acquisition are necessary to test well-formulated hypotheses regarding temporal dynamics. Besides stimulus-evoked activity (i.e., ERP), oscillatory features of scalp EEG are sensitive to the timing because its coherent coupling reflects the information flow of macroscopic neural activities (Fries 2005, 2015). For instance, stimulus presentation in a brain-state-dependent manner is one promising approach to test the input-output relationship of brain regions (Bergmann et al. 2019; Madsen et al. 2019; Stefanou et al. 2018; Zrenner et al. 2018). As the LSL-based application enables the online integration of EEG, behavioral data, and peripheral recordings, it would extend the state-dependent stimulation to the interaction of multimodal data-dependent, which potentially elucidates the mechanism underlying complex neural-body interaction (Al et al. 2020). Thereby, the LSL-based system is beneficial to minimize the effect of jitters that may blur the main effect of interest and allow accurate testing of the hypotheses. Moreover, the online acquisition of temporally aligned multimodal data would enable intuitive and robust neural interfaces, such as EEG-based brain-computer interfaces combined with eye movement or pupil responses (Mathôt et al., 2016; Joshi et al., 2020). The efficacy of online integrated analysis for the multimodal signals is of potential importance to realize accurate readout of human intention in a non-invasive manner.

## Conflict of Interest

J.U. is a founder and representative director of the university startup company, LIFESCAPES Inc. involved in the research, development, and sales of rehabilitation devices, including brain-computer interfaces. He receives a salary from LIFESCAPES Inc., and holds shares in LIFESCAPES Inc. This company does not have any relationships with the device or setup used in the current study. The remaining authors declare no competing interests.

## Author Contributions

**Seitaro Iwama**: Methodology, Software, Formal analysis, Investigation, Data curation, Writing - original draft, Writing - review & editing, Visualization. **Mitsuaki Takemi**: Conceptualization, Writing - original draft, Writing - review & editing, Visualization, Project administration, Supervision. **Ryo Eguchi**: Software, Writing - review & editing. **Ryotaro Hirose**: Data curation, Writing - review & editing. **Masumi Morishige**: Software, Investigation, Writing - review & editing. **Junichi Ushiba**: Conceptualization, Resources, Writing - review & editing, Supervision, Funding acquisition.

## Funding

This study was supported by JST Moonshot R&D Grant Number JPMJMS2012, Japan.

## Acknowledgments

We thank Shoko Tonomoto, Aya Kamiya and Yui Yoshioka for their general support.

## Notes

https://github.com/Junichi-Ushiba-Laboratory/U-BCI

